# Gremlin 1 is required for macrophage M2 polarization

**DOI:** 10.1101/2022.12.07.519468

**Authors:** Liberty Mthunzi, Simon C Rowan, Ulla G Knaus, Paul McLoughlin

## Abstract

Pro-proliferative, M2-like polarization of macrophages is a critical step in the development of fibrosis and remodeling in chronic lung diseases such as pulmonary fibrosis and pulmonary hypertension. Macrophages in healthy and diseased lungs express gremlin 1 (Grem1), a secreted glycoprotein that acts in both paracrine and autocrine manners to modulate cellular function. Increased Grem1 expression plays a central role in pulmonary fibrosis and remodelling, however, the role of Grem1 in M2-like polarization of macrophages has not previously been explored. The results reported here show that recombinant Grem1 potentiated M2-like polarization of mouse macrophages and bone marrow derived macrophages (BMDM) in response to the Th2 cytokines IL4 and IL13. Genetic depletion of Grem1 in BMDMs inhibited M2 polarization while exogenous gremlin 1 could partially rescue this effect. Taken together, these findings reveal that gremlin 1 is required for M2-like polarization of macrophages.

## Introduction

Both resident and recruited macrophages have important roles in lung inflammation, fibrosis and remodelling. Resident macrophages arise during embryogenesis and are distinct from macrophages derived from postnatal recruitment and differentiation of monocytes during acute inflammation, which originate in the bone marrow(1). These macrophages promote innate immune responses by producing inflammatory cytokines that recruit neutrophils and other leukocytes and subsequently regulate the resolution of inflammation and repair through clearance of cellular debris, interaction with the alveolar epithelium, and production of growth factors (1-5). The switch of macrophages to an alternative, pro-repair phenotype (M2) is driven by type 2 immune processes, which are characterized by increased concentrations of interleukin IL4, IL5, IL9 and IL13 (6). However, persistent M2-like macrophage activation contributes to the development of fibrosis and remodelling in chronic lung diseases such as pulmonary fibrosis and pulmonary hypertension (2, 5-8). M2-like polarization of macrophages changes the expression of multiple genes including increased expression of arginase 1 (Arg1), found in inflammatory zone 1 (Fizz1) and the mannose receptor (Mrc1) (5).

Macrophages in healthy and diseased lungs express gremlin 1 (Grem1) (9, 10). Grem1 is a secreted glycoprotein that acts both in a paracrine and autocrine fashion modulating the behaviour of cells secreting it and those in the immediate vicinity (11). Normal expression of Grem1 is required in adulthood for the regulation of cellular proliferation, differentiation and homeostatic functions including the maintenance of the normal bowel epithelium, bone marrow hematopoiesis and adipogenesis (12-14). In contrast, excessive Grem1 promotes abnormal cellular differentiation and cancer development (15, 16). Abnormally increased Grem1 also plays a pathogenetic role in the development of a number of important lung diseases including pulmonary fibrotic diseases (9, 10, 17) and pulmonary hypertension (18).

Taken together, these previous reports suggest that Grem1 may have a role in regulating macrophage function, in particular polarization towards the M2 phenotype that contributes to the development of chronic lung diseases. The aim of the experiments reported here was to examine the effects of Grem1 expressed in macrophages and Grem1 derived from other cell sources on M2-like activation of macrophages by the type 2 cytokines IL4 and IL13.

## Materials and Methods

### Materials

Anti-gremlin 1 antibody was purchased from R&D Systems and used at a final concentration of 40 ug/mL (Cat #: AF956, Lot #: GBF0513071). Anti-goat secondary Ab was purchased from Vector Labs and used at a final concentration of 4 ug/mL (Cat #: BA-5000, Lot #: DF507). RNeasy Mini Kit for RNA extraction was purchased from Qiagen (Cat #: 74014). We purchased Superscript III Reverse Transcriptase (Cat #: 18080-44) from Invitrogen for use in our cDNA synthesis reaction. IL4 (Cat #: BC1817031, Lot #: BC1716022), IL13 (Cat #: 413-ML-005, Lot #: 413-ML-005) and gremlin 1 (Cat #: 956-GR-050, Lot #: ESD4616121) recombinant proteins were all purchased from R&D Systems. Gusb, Arg1, Fizz1, Mrc1, Grem1 primer/probes were all purchased from Thermo Fisher Scientific. All primer/probes except Gusb (VIC) had FAM fluorescent dye attached.

### Mice

All protocols and procedures were approved by University College Dublin’s Animal Research Ethics Committee and licensed by the Health Products Regulatory Authority of Ireland. Male and female mice (3 – 6 months old) were used for all studies. Grem1fl^x/+^ mice (VelociGene modified allele ID number 1083) were mated with B6;129-Gt(Rosa)26Sortm2(icre/ERT2Nat)/J mice (Stock number 004847; Jax Laboratories, Maine, USA) to generate R26Cre-GREM1^fl/fl^ mice as previously described (12). Mice were housed in climate-controlled rooms under a 12h light/ dark cycle, with ad libitum access to water and food. Excision of Grem1 sequences in R26Cre-Grem1^fl/fl^ mice was induced as previously described by providing *ad libitum* access to chow with added tamoxifen (400 mg/tamoxifen citrate/kg diet, Envigo, Huntingdon, UK) for eight days. Controls were given access to standard chow for an identical period; mice were randomly allocated to each group (12). Mice were sedated by inhalation of isoflurane then anesthetized by intraperitoneal injection (I.P) of sodium pentobarbitone (70 mg/kg). After confirming depth of anesthesia by absence of response to paw compression, the femoral artery was incised, and the animal killed by exsanguination.

### Alveolar Macrophage (AM) Cell Culture

Following euthanasia, bronchoalveolar lavage (PBS, 1 mL x 5 times) was completed. The collected fluid was centrifuged (400g, 4min, 4°C) and the cell pellet re-suspended in cell culture medium (DMEM supplemented with FBS (10%, vol/vol) and HEPES (10 mM)) and allowed to settle on coverslips for two hours (37°C, 95% air, 5% CO_2_). Following this, non-adherent cells were washed away. Cells were confirmed as macrophages by characteristic morphology and expression of the macrophage markers CD68 and F4/80 (data not shown).

### Bone Marrow-Derived Macrophage Cell Culture

Bone marrow-derived macrophages (BMDMs) were generated from R26Cre-Grem1^*fl/fl*^ mice fed standard chow or chow with added tamoxifen and from wildtype C57BL6/J mice, using standard protocols (19). Briefly, following euthanasia, bone marrow was removed from the femurs and tibiae and the cells were suspended in PBS. After straining the suspension (70 µm, Fisher Scientific), red cells were lysed in lysis buffer, the remaining cells were collected by centrifugation (400g, 4min, 4°C) and resuspended in cell culture medium supplemented with L929-cell conditioned medium (LCM, 20%, vol.vol^-1^). The bone marrow progenitor cells were cultured in this medium for seven days with a change of medium on the third day. Post maturation, cells were confirmed as macrophages by adhesion to cell culture dishes, morphology and expression of macrophage markers CD68 and F4/80 (data not shown). In some experiments, Grem1 depletion was induced in macrophages *in vitro* by incubating cells from R26Cre-Grem1^*fl/fl*^ mice fed a normal diet in cell culture medium containing 4-hydroxytamoxifen (4-OHT, 2 µM).

### Polarization of macrophages in vitro

AMs and BMDMs were polarized by exposure of cells to IL4 and IL13 (20 ng/ml each) for 48h.

### Immunostaining

For immunostaining, cells on coverslips were fixed in methanol (100% vol.vol^-1^) for 10mins at 4ºC. Non-specific staining was blocked by incubating coverslips in rabbit serum diluted in PBS (1:10) at room temperature for 1h. The blocking serum was removed, and coverslips were incubated in primary antibody overnight at 4ºC. The next day, slides were washed with buffer (0.1% vol.vol^-1^ Tween in PBS), then incubated with a biotin-labelled, anti-goat IgG antibody (BA-5000, Vector Labs) at room temperature for 1h. Coverslips were then washed in buffer and incubated with streptavidin-linked Alexa fluorophore® 594 conjugate (1 µg/mL, Thermo Fisher) for 1h. Coverslips were then washed again, counterstained with DAPI (0.2 µg/mL, Sigma-Aldrich) and mounted onto slides using fluorescent mounting medium (DAKO, CA). Slides were imaged by epifluorescence microscopy (CKX41, Olympus, Japan) no downstream processing was performed.

### Analysis of mRNA expression

mRNA was extracted using TriReagent (Sigma-Aldrich, St. Louis) according to the manufacturer’s protocol along with glycoblue co-precipitant (1.6 mg/ml, Sigma-Aldrich). mRNA was re-suspended in RNase and DNase-free water. Total RNA from cells (500 – 1000 ng) was reverse transcribed to cDNA using Superscript III Reverse Transcriptase cDNA synthesis kit (Invitrogen) as per manufacturer’s protocol. Real-time PCR was performed on 384-well plates and each sample was measured in duplicate. *GusB* was used as the endogenous control (expression was unaffected by experimental conditions) according to the Taqman PCR protocol (Applied Biosystems). All primers and probes were 90 – 100% efficient as examined by standard curve analysis (data not shown). Target mRNA expression was assessed using the standard curve method and expressed relative to the mean of the control group value.

### Statistical analysis

All data are presented as median (interquartile range, IQR). For statistical analysis data were normalized by log transformation and the significance of differences between group means was determined using paired or unpaired t tests as appropriate. For statistical analysis of non-parametric data, we used the Wilcoxon signed-rank test or Mann-Whitney U as appropriate. Correction for multiple comparisons of means where required was undertaken using the Holm-Sidak step down method (20). A P value < 0.05 was considered statistically significant. Where P values are >0.001, the exact value is shown. All analyses were undertaken using the statistical package for the social sciences (SPSS, Version 28, IBM).

### Source Data

Source data for all figures can be accessed using the publicly available DOI for Figshare data: https://doi.org/10.6084/m9.figshare.21158914

## Results

### Alveolar macrophages and bone marrow derived macrophages express Gremlin 1

Immunostaining of AMs isolated from healthy, wild type mice (n=6) using anti-Grem1 antibody demonstrated positive staining (Figure 1A), confirming previous reports that AMs express Grem1 (9, 10). No staining was seen when primary antibody was omitted or when a similarly produced rabbit polyclonal control antibody (goat anti-HAND) was used (data not shown). Stimulation of AMs *in vitro* with the type 2 cytokines IL4 and IL13 caused increased expression of the classic M2 polarization markers Arg1 and Fizz1, although Mrc1 was unchanged (Figure 1C-E). Interestingly, M2 polarization did not alter Grem1 mRNA expression (Figure 1F).

**Figure 1.**
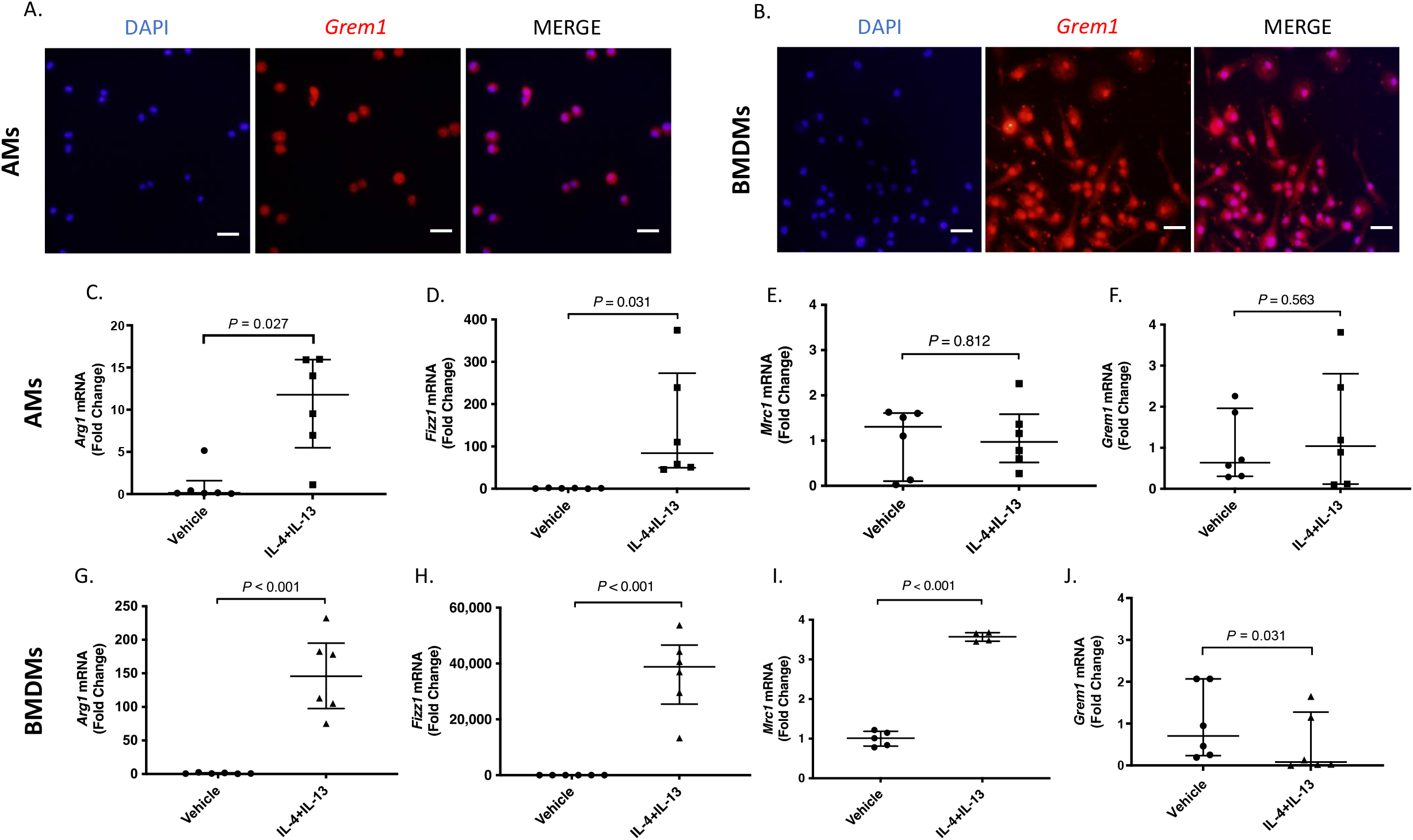
Immunostaining of gremlin 1 (red) shows expression in alveolar macrophages (A) and bone marrow derived macrophages (B) in representative images from six mice. Effects of exposure of alveolar macrophages (AMs) to IL4 and IL13 on the expression of the M2-like markers, *Arg1* (C), *Fizz1* (D) *Mrc1* (E) and *gremlin* 1 (F). Effects of exposure of bone marrow derived macrophages (BMDMs) to IL4 and IL13 on the expression of *Arg1* (G), *Fizz1* (H), *Mrc1* (I) and *gremlin* 1 (J). The scale bar indicates 20 μm (x40 objective, NA 0.9).

Grem1 expression was detected in BMDMs basally (M0 polarization) by immunostaining (Figure 1B). Specificity of the anit-Grem1 antibody has previously been demonstrated by both siRNA directed against Grem1 and demonstration of blocking of Grem1 function (18). Stimulation of BMDMs *in vitro* by IL4/IL13 produced a change in morphology to an elongated phenotype (data not shown) and an increase in the M2 polarization markers Arg1, Fizz1 and Mrc1 (Figure 1G, H, I). Interestingly, Grem1 expression was reduced following M2 polarization (Figure 1J).

### Grem1 is required for polarization of macrophages to an M2-like phenotype

We next examined the effect of recombinant human (rh)Grem1 on the polarization response of BMDMs from wild type mice. IL4 and IL13 stimulation caused increased expression of Arg1, Fizz1 and Mrc1, as previously (Figure 2A, B and C). Addition of rhGrem1 to IL4 and IL13 did not further change Mrc1 expression but caused a significant further increase in the expression of Arg1 and Fizz1 (Figure 2A-C), when compared to IL4 and IL13 alone, suggesting that Grem1 produced by cellular sources adjacent to macrophages might modulate polarization. Grem1 alone did not alter Arg1, Fizz1 or Mrc1 expression (data not shown).

**Figure 2.**
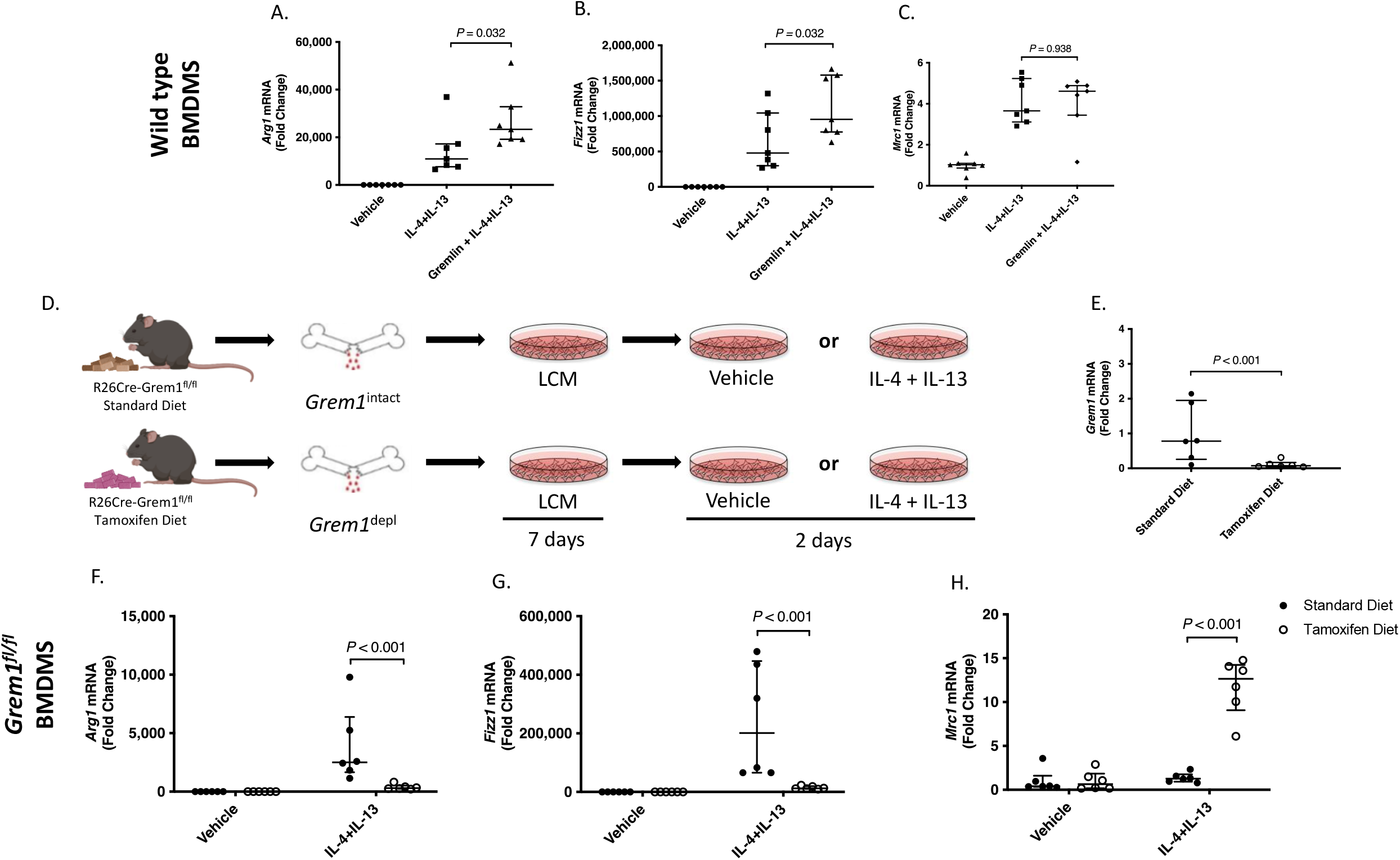
Effect of Gremlin 1 on *Arg1* (A), *Fizz1* (B) and *Mrc1* (C) expression in BMDMs isolated from wild type mice. Schematic (D) showing experimental protocol to examine the effect of *in vivo* depletion of gremlin 1 on the subsequent responses to IL4 and IL13. Effect of tamoxifen on Grem1 expression (E). Effect IL4 and IL13 alone or together with rhGremlin1 on the expression of *Arg1* (F), *Fizz1* (G) and *Mrc1* (H) in BMDMs from Grem1^intact^ mice (standard chow, filled circles) and from Grem1^depl^ mice (tamoxifen chow, open circles).

To examine the effects of Grem1 produced by the macrophages on their own polarization, we used BMDMs obtained from R26Cre-Grem1^fl/fl^ mice fed either standard chow (Grem1^intact^) or tamoxifen containing chow (Grem1^depl^). At the time of euthanasia, the mice showed normal weight and behaviour. LCM-differentiated BMDMs from both groups were stimulated with IL4/IL13 or exposed to vehicle alone (protocol schematically illustrated in Figure 2D). In mice that received tamoxifen chow, Grem1 was successfully depleted (Figure 2E) when compared to those receiving a standard diet. Grem1 depletion caused marked changes in the polarization response to IL4/IL13, significantly reducing Arg1 and Fizz1 and augmenting Mrc1 expression (Figure 2F, G and H). BMDMs isolated from wild type mice fed tamoxifen and exposed to IL4/IL13 showed similar M2-like polarization responses to those of wild type mice fed a normal diet (data not shown).

### Exogenous Grem1 partially restores the polarization responses of Grem1^depl^ macrophage

We have previously shown that more prolonged tamoxifen administration (>12 days) to induce Grem1 depletion in R26Cre-Grem1^fl/fl^ mice causes bowel abnormalities, bone marrow failure and is lethal (12). Although such mice fed tamoxifen typically did not become unwell after eight days of tamoxifen exposure (and none in the study presented here), there remained the possibility that the abnormal polarization responses observed following *in vivo* depletion of Grem1 were, at least in part, due to the effects of an abnormal bone marrow environment caused by Grem1 loss that persisted *in vitro* (12). For this reason, we also examined the effect of *in vitro* depletion of Grem1 during cell culture after bone marrow harvesting. R26Cre-Grem1^fl/fl^ mice fed a normal diet were euthanised, their bone marrow isolated and the bone marrow progenitor cells then cultured with LCM to stimulate macrophage differentiation either in the presence or absence of tamoxifen (schematic illustration of protocol in Figure 3A). This caused marked depletion of Grem1 (Figure 3B), similar to that following depletion of Grem1 *in vivo* (Figure 2E). The responses of these BMDMs to IL4/IL13 were very similar to those of BMDMs following *in vivo* depletion i.e. reduced Arg1 and Fizz1 expression and increased Mrc1 expression compared to Grem1^intact^ BMDMs (Figure 3C-E). These results suggested that the altered polarization response of macrophages from mice in which Grem1 had been depleted *in vivo* was not due to any deleterious effect of Grem1 loss on the bone marrow but due to the loss of Grem1 produced by the macrophages.

**Figure 3.**
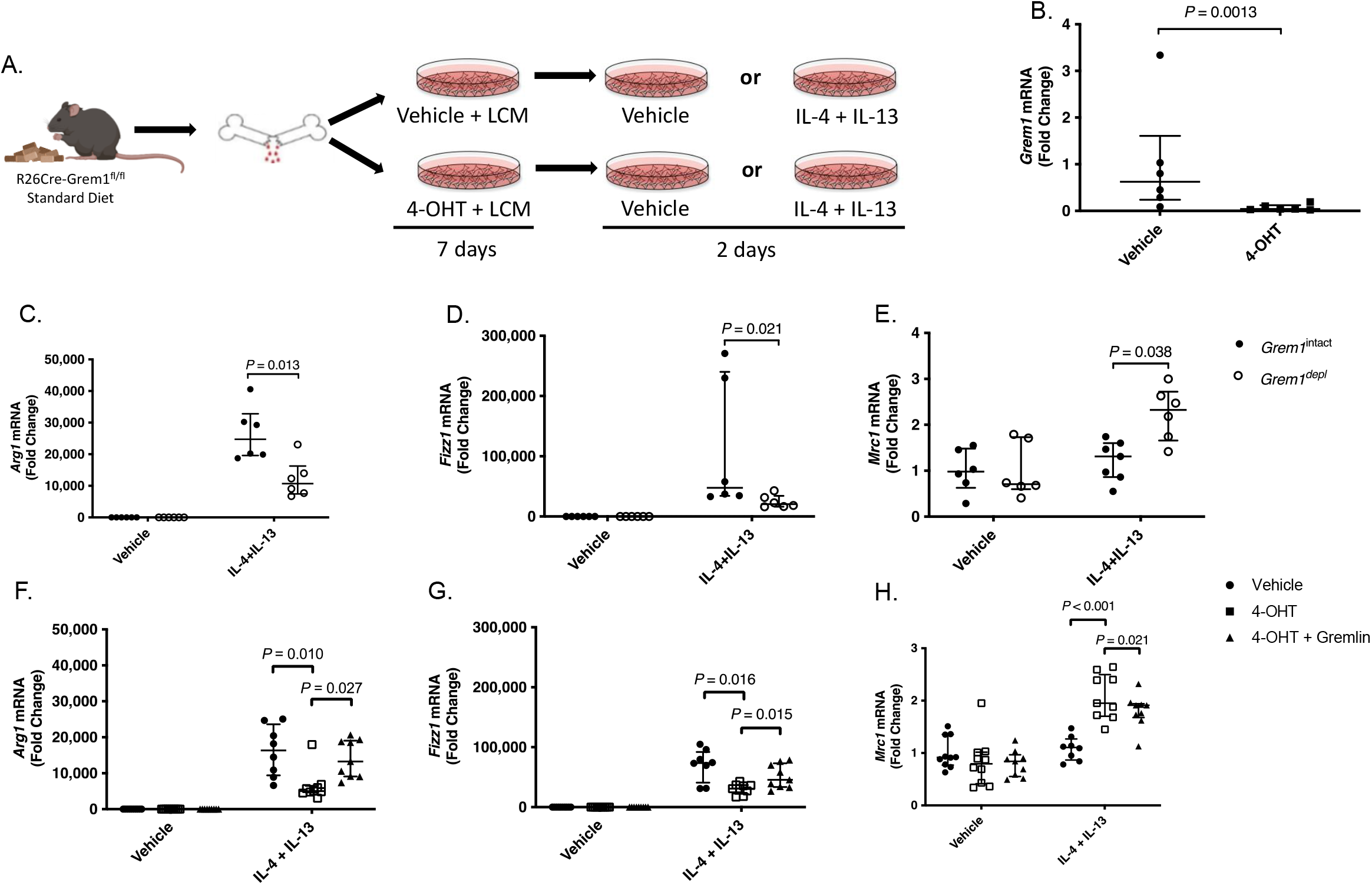
Schematic showing experimental protocol to examine the effect of *in vitro* depletion of gremlin 1 on the subsequent responses to IL4 and IL13 (A). Addition of tamoxifen (4OHT) *in vitro* reduced Grem1 expression (B). Effect of IL4/IL13 on the expression of *Arg1* (C), *Fizz1* (D) and *Mrc1* (E) in Grem1 intact bone marrow derived macrophages. Effect of IL4/IL13 either alone or together with rhGremlin 1 on the expression of *Arg1* (F), *Fizz1* (G) and *Arg1* (H) in *Grem1* depleted bone marrow derived macrophages.

Since exogenous Grem1 potentiated wild type macrophage activation (Figure 1A-C), we hypothesized that exogenous Grem1 might restore the M2 polarization responses of macrophages depleted of endogenous Grem1. When rhGrem1 was added to BMDMs that had been depleted of Grem1 *in vitro*, the reduced Arg1 and Fizz1 expression observed in Grem1^depl^ BMDMs was increased towards the values observed in Grem^intact^ BMDMs (Figure 3F and G). The increased *Mrc1* expression induced by IL4/IL13 in Grem1^depl^ BMDMs was significantly reduced by the addition of rhGrem1 but not restored to that seen in the Grem1^intact^ BMDMs (Figure 3H). Exposure of BMDMs derived from wild type mice to tamoxifen in the same way did not alter IL4/IL13 driven polarization (Figure 4). We conclude from these data that exogenously administered Grem1 can partially restore macrophage responses to IL4/IL13 stimulation.

**Figure 4.**
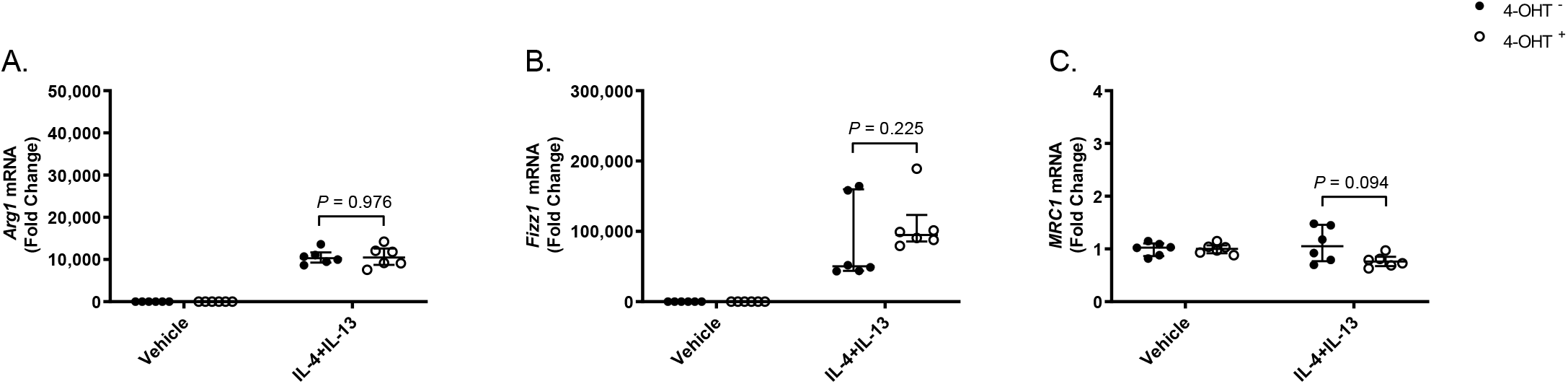
Addition of tamoxifen (4-OHT) *in vitro* did not change effect of IL4 and IL13 on wildtype BMDM polarization. Effect 4-OHT on IL4 and IL13 induced expression of *Arg1, Fizz1* and *Mrc1* in wildtype bone marrow derived macrophages (A, B, C).

## Discussion

These results show that gremlin 1 is required for M2-like polarization of macrophages in response to the Th2 cytokines IL4 and IL13. We report that exogenous Grem1 potentiates the polarization of macrophages as characterized by increased expression of the pro-fibrotic M2 polarization markers Arg1 and Fizz1. Depletion of endogenous Grem1 impaired macrophage polarization but exogenous administration of Grem1 could partially restore the M2 polarization of Grem1-depleted macrophages.

It is now well recognised the M1 and M2 polarization of macrophages represent two ends of a spectrum and that expression of many genes and proteins are changed along this spectrum. We examined expression of Mrc1 as it is a canonical marker of M2-like polarization. We also examined Arg1 and Fizz1 because, in addition to being well recognised markers of M2-like polarization, they also play important roles in the mechanisms underlying pulmonary fibrosis and vascular remodelling (21, 22). Thus, our findings suggest that increased Grem1 potentiates the profibrotic actions of macrophages in fibrotic lung disease.

The effects of Grem1 that we observed might have been mediated via a number of different signalling pathways. Grem1 was first identified as an antagonist of bone morphogenetic proteins 2, 4 and 7 but was subsequently shown to bind to VEGFR2, SLITs, macrophage migration inhibitory factor (MIF) and FGF21 resulting in altered signalling through these pathways (11, 23-27). Grem1 from both autocrine and paracrine sources could act though these pathways. Recent scRNAseq studies have shown that cells known to interact with macrophages express Grem1 in the fibrotic lung including fibroblasts, myofibroblasts and alveolar epithelial cells (28).

Grem1 can also act through intracellular actions suggesting that Grem1 produced by adjacent cells may not be able to completely correct the changes in polarization caused by genetic depletion of Grem1 within the macrophages. Further work will be required to determine through which of these mechanisms Grem1 acts on macrophage polarization.

Macrophage polarization toward an M2 phenotype is an important step in a number of fibro-proliferative diseases including pulmonary fibrosis and pulmonary hypertension. We have discovered a previously unknown role for Grem1 in the regulation of macrophage polarization that may represent a novel cellular mechanism by which increased Grem1 promotes fibrosis and remodelling in lung disease.

